# Tornadic shear stress induces a transient, calcineurin-dependent hyper-virulent phenotype in Mucorales molds

**DOI:** 10.1101/2020.02.14.945998

**Authors:** Sebastian Wurster, Alexander M. Tatara, Nathaniel D. Albert, Ashraf S. Ibrahim, Joseph Heitman, Soo Chan Lee, Amol C. Shetty, Carrie McCracken, Karen T. Graf, Antonios G. Mikos, Vincent M. Bruno, Dimitrios P. Kontoyiannis

**Author notes:** Corresponding author: Prof. Dimitrios P. Kontoyiannis, MD, ScD, PhD (hon), The University of Texas M.D. Anderson Cancer Center, Department of Infectious Diseases, Infection Control and Employee Health, 1515 Holcombe Boulevard, Unit 1460, Houston, TX, 77030, United States of America.

## Abstract

Trauma-related necrotizing myocutaneous mucormycosis (NMM) has a high morbidity and mortality in victims of combat-related injuries, geo-meteorological disasters, and severe burns. Inspired by the observation that several recent clusters of NMM have been associated with extreme mechanical forces (e.g. during tornados), we studied the impact of mechanical stress on Mucoralean biology and virulence in a *Drosophila melanogaster* infection model. In contrast to other experimental procedures to exert mechanical stress, tornadic shear challenge (TSC) by magnetic stirring induced a hyper-virulent phenotype in several clinically relevant Mucorales species but not in *Aspergillus* or *Fusarium*. Whereas fungal growth rates, morphogenesis, and susceptibility to noxious environments or phagocytes were not altered by TSC, soluble factors released in the supernatant of shear-challenged *R. arrhizus* spores rendered static spores hyper-virulent. Consistent with a rapid decay of TSC-induced hyper-virulence, minimal transcriptional changes were revealed by comparative RNA sequencing analysis of static and shear-challenged *Rhizopus arrhizus*. However, inhibition of the calcineurin/heat shock protein 90 (hsp90) stress response circuitry by cyclosporine A (CsA) and tanespimycin abrogated the increased pathogenicity of *R. arrhizus* spores following TSC. Similarly, calcineurin loss-of-function mutants of *Mucor circinelloides* displayed no increased virulence capacity in flies after undergoing TSC. Collectively, these results establish that TSC induces hyper-virulence specifically in Mucorales and point out the calcineurin/hsp90 pathway as a key orchestrator of this phenotype. Our findings invite future studies of topical calcineurin inhibitor treatment of wounds as an adjunct mitigation strategy for NMM following high-energy trauma.

**Significance:** Given the limited efficacy of current medical treatments in trauma-related necrotizing mucormycosis, there is a dire need to better understand the Mucoralean pathophysiology in order to develop novel strategies to counteract fungal tissue invasion following severe trauma. Here, we describe that tornadic shear stress challenge transiently induces a hyper-virulent phenotype in various pathogenic Mucorales species but not in other molds known to cause wound infections. Our data support a model whereby shear stress-induced hyper-virulence is primarily driven by soluble factors and orchestrated by the calcineurin/hsp90 pathway. Importantly, pharmacological and genetic inhibition of calcineurin signaling abrogated hyper-virulence in shear stress-challenged Mucorales, encouraging further evaluation of (topical) calcineurin inhibitors to improve therapeutic outcomes of NMM after combat-related blast injuries or violent storms.

## Introduction

Necrotizing myocutaneous invasive mold infections following severe trauma represent a life-threatening disease with high morbidity and mortality (1–4). A variety of molds have been implicated as causative agents, with fungi belonging to the order Mucorales predominating as the most common and devastating cause (3–4). These emerging fungal pathogens affect an expanding population of hosts and are characterized by innate resistance to many antifungals, broad geographic and environmental distribution, and high virulence (5–8). While immunocompromised patients are at highest risk for the development of mucormycosis (5–8), immunocompetent individuals are prone to necrotizing myocutaneous mucormycosis (NMM) when incurring penetrating trauma (2, 9–12), combat-related wounds (13–15), or burn injuries (1).

Interestingly, several clusters of NMM have been observed after trauma events in settings of extreme mechanical forces. Specifically, NMM has emerged in military personnel suffering wound infections after blast injuries from improvised explosive devices in Afghanistan and Iraq, and the recovery of Mucorales from wound cultures was frequently associated with recurrent tissue necrosis (15). Moreover, a cluster of NMM cases due to *Apophysomyces trapeziformis*, an uncommon agent of mucormycosis, was reported after the 2011 EF-5 Joplin tornado, causing considerable mortality and necessitating aggressive debridement and complex reconstruction among survivors (10). NMM cases were also reported after the 2004 Indian Ocean tsunami (9, 16). Given that NMM is encountered after high-energy events, we hypothesized that mechanical stress challenge may alter the virulence traits and eventually result in increased pathogenicity of Mucorales molds.

Studies in bacteriology suggest that prokaryotic pathogens experience and sense a variety of mechanical events including shear forces that can modulate motility, surface adhesion, and biofilm formation (17–18). There is increasing evidence that shear forces also influence eukaryotic cell behavior, proliferation, and signaling (19–20), and previous reports have highlighted that cascades involved in environmental stress response serve crucial roles in controlling fungal morphogenesis and pathogenic capacity (21–22). However, the specific link between mechanical stress and fungal virulence is as yet uncharacterized. To address this gap of knowledge in the context of trauma-related NMM, we studied the impact of tornadic shear forces on the morphogenesis, transcriptional signatures, and pathogenicity of Mucorales. Using a *Drosophila melanogaster* mucormycosis model that has been validated to recapitulate key virulence attributes of human infections (23–24), we found a transient, hyper-virulent phenotype in shear-challenged Mucorales but not in other common opportunistic molds. Our findings further reveal that shear force-induced hyper-virulence in Mucorales relies on the calcineurin/heat shock protein 90 (hsp90) pathway, giving rise to new potential avenues of therapeutic interventions in NMM following combat injuries or geo-meteorological disasters.

## Results

### Tornadic shear challenge induces a unique, transient, hyper-virulent phenotype of Mucorales

At first, we compared the influence of different tornadic shear challenge (TSC) procedures on the *in vivo* pathogenicity of *Rhizopus arrhizus*, the most common cause of mucormycosis (25). Spore suspensions were either centrifuged, vortexed, or stirred with a magnetic stir rod for 30 min. Wild-type *Drosophila melanogaster* flies infected with static *R. arrhizus* spores showed 7-day survival rates of 37-41 % (**Fig. S1A-C**), consistent with our previous findings (23, 26). Neither centrifuged (**Fig. S1A**) nor vortexed (**Fig. S1B**) spores elicited increased mortality in infected flies compared with static spores (p = 0.89 and 0.91, respectively). In contrast, magnetic stirring of *R. arrhizus* spores led to near universal lethality of flies in 7 days (6 % survival versus 41 %, p < 0.001) and reduced the median survival time of infected flies from 5 to 2 days (**Fig. S1C**), suggesting enhanced fungal pathogenicity. Therefore, magnetic stirring was used to simulate TSC in all subsequent experiments.

In order to evaluate the generalizability of TSC-induced hyper-virulence in Mucorales, we tested additional clinical isolates of *R. arrhizus, Rhizomucor pusillus*, and *Mucor circinelloides*. For all three strains, TSC led to a precipitous decline in 7-day survival rates of infected WT flies from 36-44 % to 8-14 % (p < 0.001, **Fig. 1A**). Interestingly, the increase in virulence after TSC was particularly pronounced in *A. trapeziformis*, with a 26 % 7-day survival rate of infected flies compared with 64 % for flies infected with static spores (p < 0.001, **Fig. 1A**), concordant with the dominance of *A. trapeziformis* after high-energy trauma (1, 10, 27). However, as it is challenging to obtain large quantities of *Apophysomyces* spores, we used *R. arrhizus* as a model organism for further investigations of TSC-induced pathogenicity. In contrast to Mucorales, the pathogenicity of opportunistic Ascomycetes molds *Aspergillus fumigatus* and *Fusarium solani* was not altered by magnetic stirring (**Fig. 1B**), indicating that TSC induces increased virulence specifically in Mucorales.

**Figure 1.**
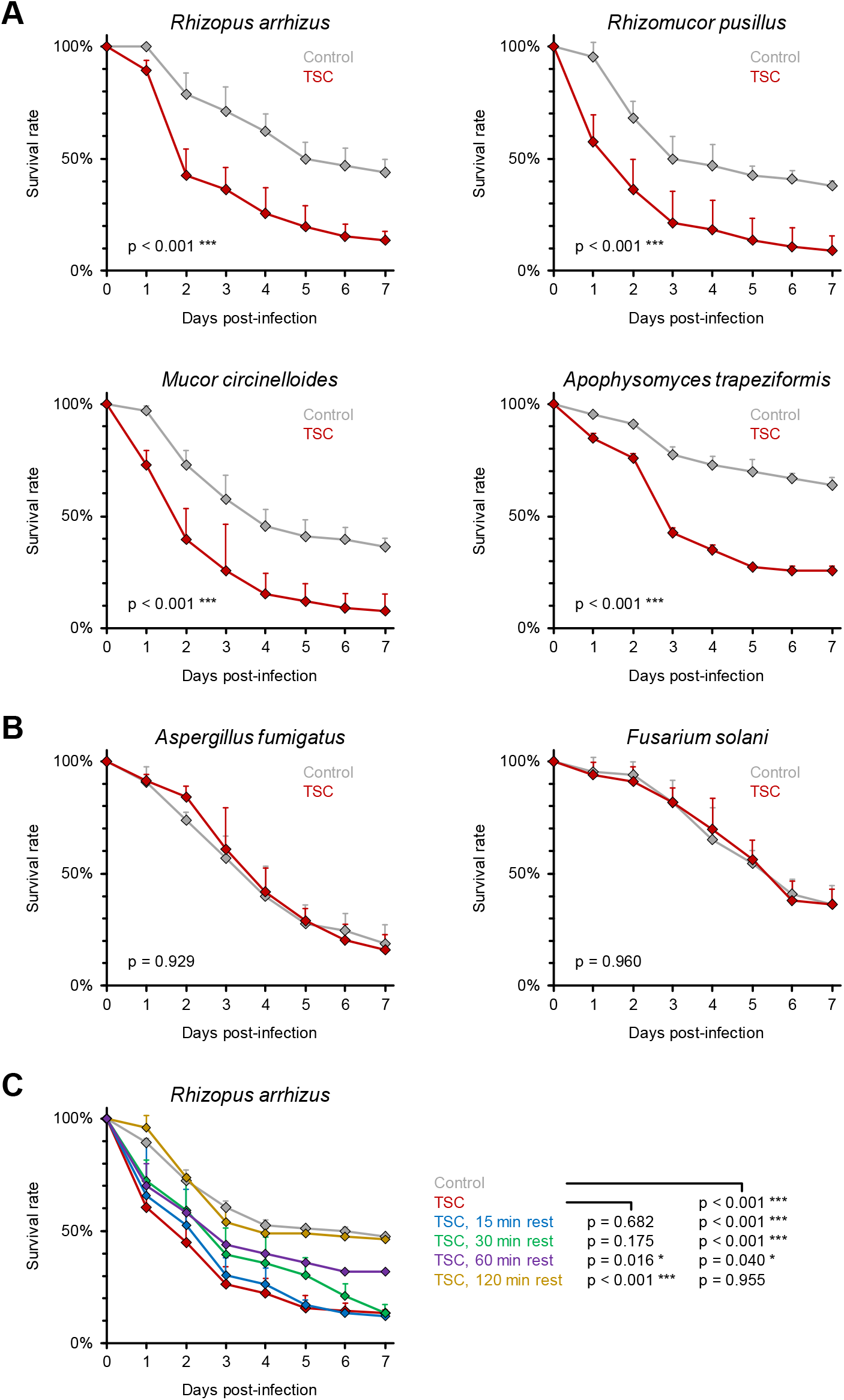
Tornadic shear challenge transiently increases the pathogenicity of Mucorales but not Ascomycetes. Legend on following page **(A-B)** Spore suspensions (10^7^/ml) of Mucorales isolates *R. arrhizus* Ra-749, *R. pusillus* Rp-449, *M. circinelloides* Mc-518*, and A. trapeziformis* CBS 125534 **(A)** as well as Ascomycetes isolates *A. fumigatus* Af-293 and *F. solani* Fs-001 **(B)** *w*ere subjected to TSC by magnetic stirring for 30 min or kept in static culture (Control). *D. melanogaster* flies were infected by pricking with a needle dipped into the spore solutions. WT flies were used for Mucorales and *F. solani*, whereas *A. fumigatus* infections were performed in *Tl^r632^/Tl^I-RXA^* mutant flies since *Aspergillus* infections are non-lethal in WT flies. For each pathogen, three independent experiments were performed with a total of 65-69 flies per condition. **(C)** WT flies were infected with *R. arrhizus* Ra-749 (10^7^/ml) either immediately after termination of TSC or after a 15-120 min resting period. The non-challenged control was inoculated within 5 minutes of the non-rested TSC cohort. Three independent experiments were performed with a total of 76 flies per condition. For all panels, survival curves were compiled from aggregated results. Error bars represent inter-replicate standard deviations. Log-rank-test.

To evaluate whether increased virulence following TSC was dependent on the infecting spore concentration, we pricked flies with a needle immersed in a range of different *R. arrhizus* spore inocula (10^4^, 10^6^, or 10^8^/ml) that underwent TSC by magnetic stirring or remained under static conditions. Expectedly, baseline mortality without TSC was inoculum-dependent, with 7-day-survival rates of 49 % (10^4^ spores/ml), 39 % (10^6^/ml), and 36 % (10^8^/ml), respectively (**Fig. S2**). For all inocula tested, *Rhizopus* infection following TSC led to significantly elevated fly mortality (p < 0.001), with essentially identical deltas in 7-day mortality rates (28-31 %). Collectively, these data suggest that enhanced pathogenicity of Mucorales after TSC is a strain-, species-, and inoculum-independent phenomenon.

We then assessed the durability of the hyper-virulent phenotype after TSC exposure of *R. arrhizus*. In line with our earlier results (**Fig. 1A**), flies pricked immediately after magnetic stirring of the fungal inoculum suspension exhibited significantly increased 7-day mortality compared with the static control (87 % versus 53 %, p < 0.001, **Fig. 1C**). However, enhanced pathogenicity in flies decayed rapidly with increasing resting periods after TSC. A 60 min resting step following TSC exposure of *R. arrhizus* spores significantly reduced fly mortality and the survival curves reverted back to the non-TSC control after 120 min post-TSC resting (**Fig. 1C**), indicating that the increased pathogenicity of Mucorales after exposure to TSC is transient.

### TSC-induced hyper-virulence of Mucorales is largely driven by secreted factors

To understand the molecular mechanisms underlying the increased virulence following TSC, we performed RNA sequencing (RNA-seq) on *R. arrhizus* spores that were either kept in static culture, exposed to TSC, or exposed to TSC and then allowed to rest for 120 minutes. We defined differentially expressed genes as those with a false discovery rate (FDR) <0.05 between experimental groups. Despite the significant sequencing depth coverage that we obtained (38.1 ± 3.62 million reads per sample), we observed that *R. arrhizus* mounted a minimal transcriptional response to TSC (**Fig. 2A**) and also to the rest following TSC (**Fig. 2B**). In fact, only 3 genes were differentially expressed between shear-challenged and static spores. Furthermore, only 22 genes were differentially expressed between the spores that were exposed to TSC and those that were allowed to rest for 120 minutes following shear challenge. All of the differentially expressed genes are uncharacterized and annotated as hypothetical proteins. Taken together, these results suggest that *R. arrhizus* does not mount a robust transcriptional response to TSC and that the increased virulence is likely not the result of transcriptional up-regulation of virulence genes.

**Figure 2.**
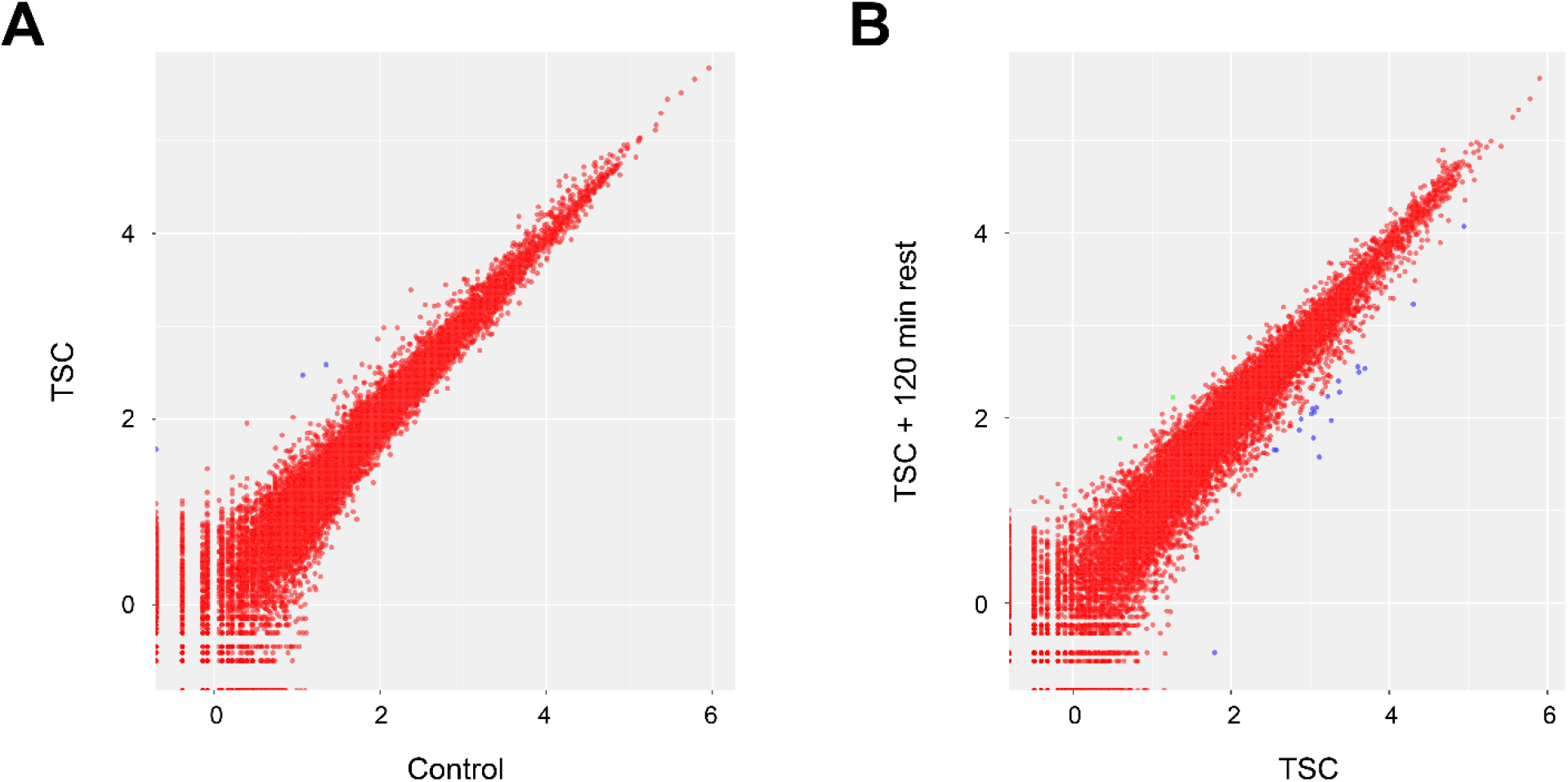
Tornadic shear challenge does not significantly alter the transcriptome of *R. arrhizus*. Scatter plots comparing the expression of each gene in *R. arrhizus* Ra-749 spores exposed to TSC or not (Control) (**A**) as well as spores exposed to TSC and then allowed to rest for 120 minutes or not (**B**). Values represent the log-transformed mean-normalized read count for each gene. Red dots indicate an FDR >0.05 (deemed not differentially expressed). Blue or green dots indicate differentially expressed genes (FDR <0.05).

Therefore, we conducted an array of phenotypic assays to further understand the features of TSC-induced hyper-virulence. Employing the recently adapted IncuCyte NeuroTrack time-lapse imaging approach (28), we tested whether TSC results in accelerated growth or hyphal filamentation of Mucorales. However, mycelial morphology (**Fig. S3A**), confluence, hyphal length, and branch point numbers (**Fig. S3B** and data not shown) remained unaffected by TSC in three different Mucorales isolates, indicating that hyper-virulence is not a result of altered mycelial expansion and/or morphogenesis.

As previous studies suggested that fungal stress adaptation can induce resistance to subsequent stress events (21–22), we further used the NeuroTrack assay to determine the impact of TSC on the Mucoralean tolerance of oxidative stress (peroxide, H_2_O_2_). In all three species tested, shear-challenged spores did not have increased resistance to subsequent peroxide exposure compared with static controls (**Fig. S4A**). Similarly, no significantly different hyphal length endpoints were found between shear-challenged and static Mucorales spores subsequently exposed to sub-inhibitory peroxide concentrations (**Fig. S4A**). In addition, simultaneous exposure to TSC and the highest sub-inhibitory peroxide concentration (1 mM) had no change on mycelial expansion compared with Mucorales spores exposed to 1 mM H_2_O_2_ in static culture (data not shown). To corroborate these data with a different stressor, we exposed shear-challenged and control spores to serial dilutions of amphotericin B and posaconazole, resulting in non-differential minimum inhibitory concentrations (MICs, **Fig. S4B**). Collectively, these data suggest that TSC does not affect the ability of Mucorales to survive and form expansive mycelium in noxious *in vitro* environments.

As increased resistance to or interference with the host’s phagocytic capacity could present another virulence feature contributing to TSC-induced pathogenicity, we next compared the phagocytic activity of *D. melanogaster* S2 hemocytes against shear-challenged and static *R. arrhizus*. S2 cells have considerable similarities with human phagocytic cells and have been previously validated as an *in vitro* system to study cellular immune responses against Mucorales (23). However, co-cultures analyzed by IncuCyte NeuroTrack imaging revealed no differential susceptibility of static and shear-challenged *R. arrhizus* spores to S2 phagocytes as indicated by comparable mycelial expansion and branching kinetics in the presence of phagocytes, regardless of prior TSC (**Fig. S5A**). In addition, invasion of host epithelia is regarded as an essential virulence trait in the establishment and progression of mucormycosis (29–30). Co-culturing *Rhizopus* spores with A549 epithelial cells for 24 and 48 h, static and TSC-exposed spores elicited comparable and inoculum-dependent release of LDH, a surrogate marker for epithelial cell damage (**Fig. S5B**).

Next, we focused our attention on the possibility that secreted metabolites could contribute to increased pathogenicity in the TSC setting. Interestingly, static *R. arrhizus* spores re-suspended in supernatants from TSC-exposed spores became equally hyper-virulent in the fly model as the TSC inoculum, with 7-day survival rates of 22 % versus 50 % with non-challenged control spores (p < 0.001, **Fig. 3A**). Inversely, the mortality of flies infected with a mixture of supernatants from static *R. arrhizus* spores and the TSC-exposed spore pellet was comparable to the unchallenged control and significantly lower than in flies infected with stirred spores (p < 0.001, **Fig. 3B**). Together, these data suggest that increased virulence of TSC-exposed spores is largely due to soluble factors.

**Figure 3.**
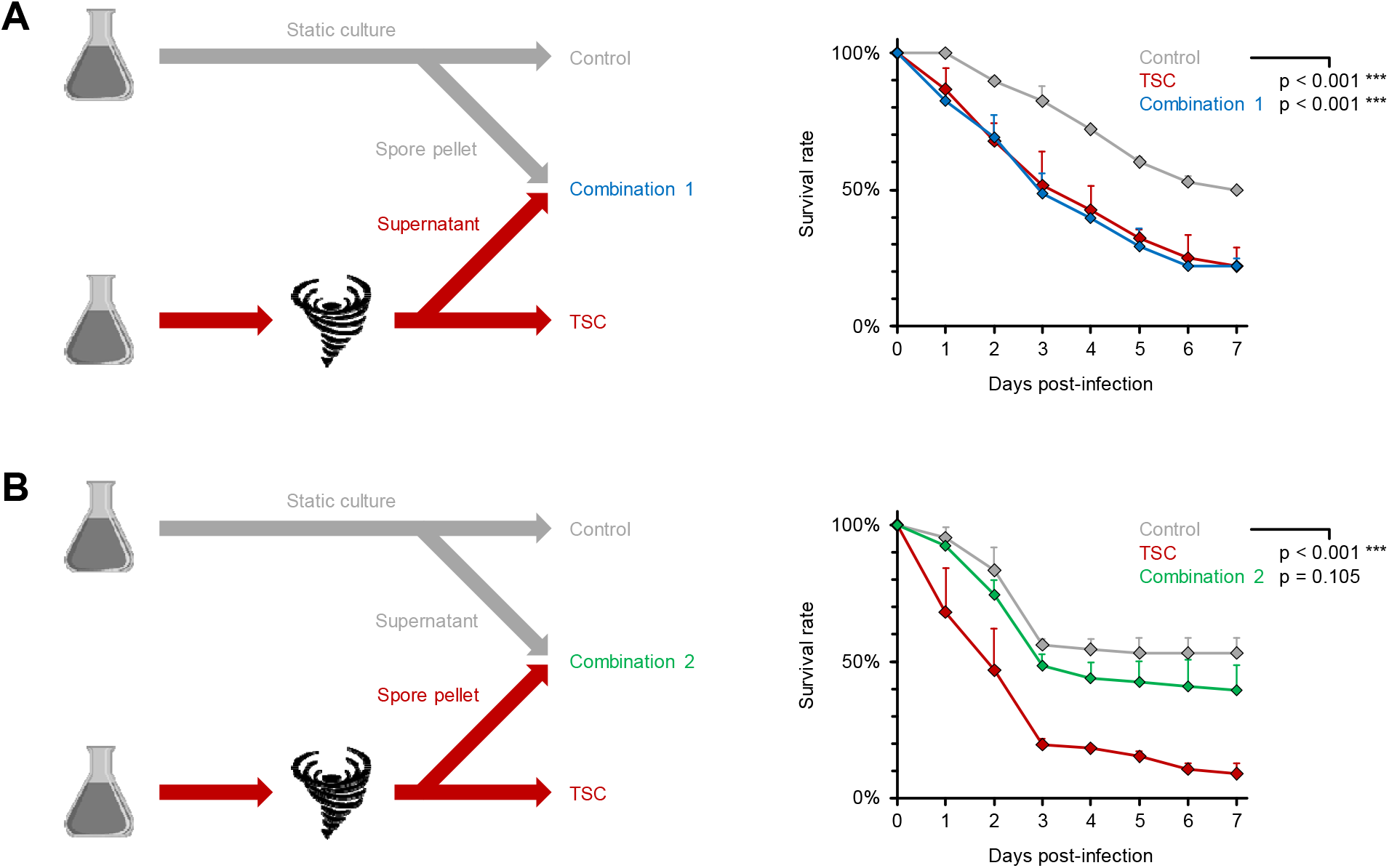
The hyper-virulent phenotype of shear-challenged Mucorales is driven by a soluble factor. *R. arrhizus* Ra-749 spore suspensions (10^7^/ml) were exposed to TSC by magnetic stirring or kept in static culture (Control) for 30 min. Additional suspensions (Combination 1 and 2) were prepared by replacing the supernatant from control spores with an equivalent volume of supernatant from TSC-exposed samples (**A**) or *vice versa* (**B**). Survival rates of infected flies were monitored for 7 days. 66-68 flies per condition were tested in three independent replicates. Error bars represent inter-replicate standard deviations. Log-rank-test.

### TSC-induced hyper-virulence depends on the Mucoralean calcineurin/hsp90 pathway

Lastly, we sought to identify cascades governing the hyper-virulent phenotype of Mucorales after TSC. As the calcineurin/hsp90 axis has been described as a pivotal driver of fungal adaptation to environmental stress (20, 22, 31–32), we hypothesized that inhibitors of this pathway may attenuate the impact of TSC on Mucoralean hyper-virulence. Indeed, addition of sub-inhibitory concentrations (100 μg/ml) of the calcineurin inhibitor cyclosporine A CsA during TSC exposure of *R. arrhizus* reduced the 7-day mortality of infected flies from 99 % to 71 % (**Fig. 4A**, p < 0.001), whereas the pathogenicity of static spores was not influenced by CsA. Furthermore, the hsp90 inhibitor tanespimycin (50 μg/ml 17-AAG, **Fig. 4B**) and its combination with CsA (**Fig. 4C**) fully reverted the pathogenicity of TSC-exposed spores to the level of the unchallenged control, further underscoring a role of the calcineurin/hsp90 pathway in TSC-induced *in vivo* pathogenicity. Importantly, CsA continued to mitigate TSC-induced hyper-virulence of *R. arrhizus* even when added after the stirring procedure (**Fig. 4D**, p < 0.001), an important finding that indicates potential feasibility of pharmacological interventions to counteract fungal virulence in NMM after high-energy events.

**Figure 4.**
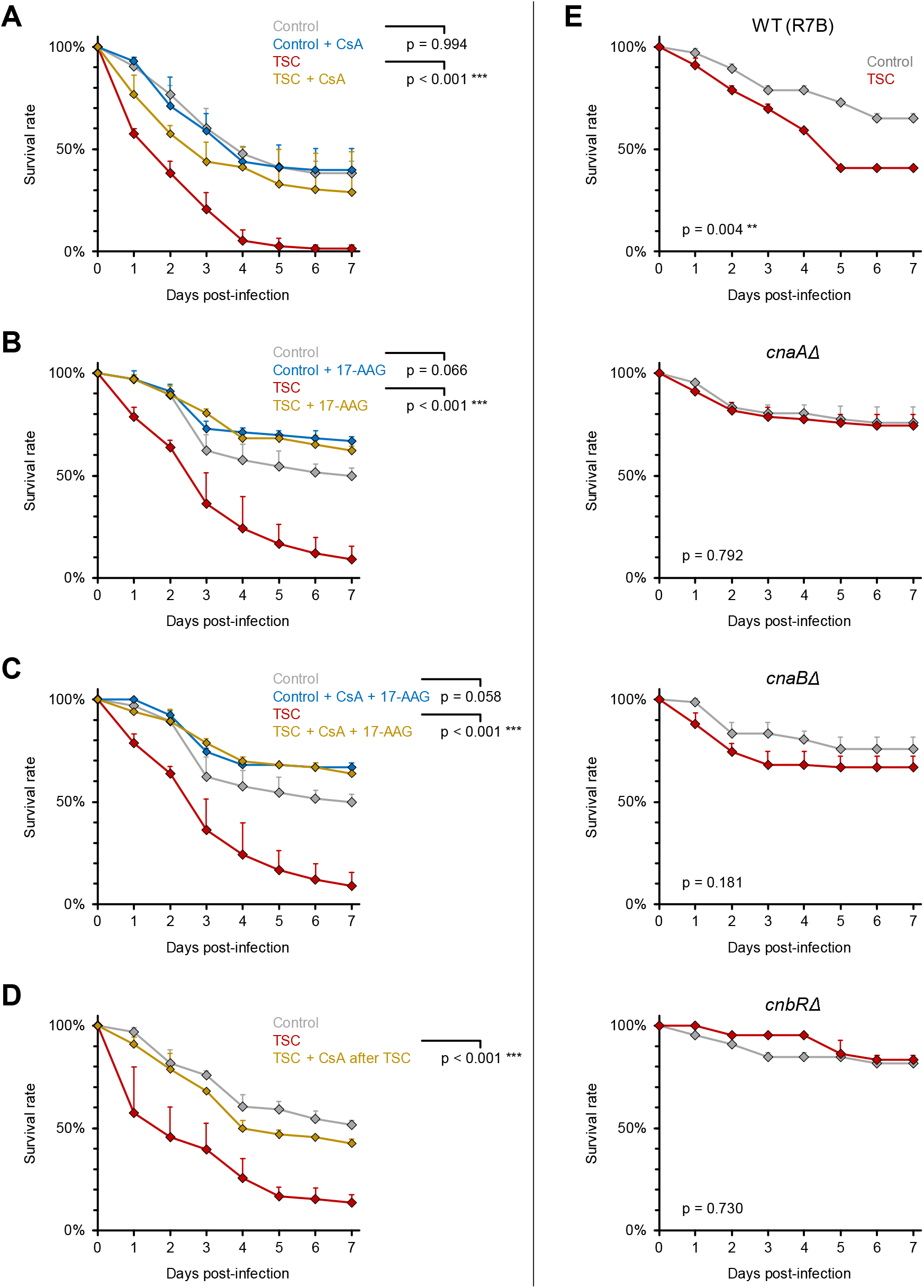
Mucoralean hyper-virulence after tornadic shear challenge depends on calcineurin and hsp90. Legend on following page (**A-C**) *R. arrhizus* Ra-749 spore suspensions (10^7^/ml) were either prepared in PBS or in PBS supplemented with inhibitors of the calcineurin/hsp90 pathway (**A**: 100 μg/ml CsA, **B**: 50 μg/ml 17-AAG, **C**: 100 μg/ml CsA + 50 μg/ml 17-AAG). Spore suspensions were exposed to TSC by magnetic stirring for 30 min or kept in static culture for the same period (Control). WT *D. melanogaster* flies were infected by pricking with a needle dipped into the spore solutions. (**D**) Flies were infected with static or shear-challenged *R. arrhizus* Ra-749 spore suspensions supplemented with 100 μg/ml CsA after the TSC procedure or not. (**E**) Flies were infected with spore suspensions of *M. circinelloides* calcineurin subunit loss-of function mutants MSL9 (*cnaAΔ*), MSL22 (*cnaBΔ*), MSL8 (*cnbR/Δ*), and the isogenic R7B control strain. For each panel, three independent experiments were performed with a total of 66-73 flies per condition. Survival curves were compiled from aggregated results. Error bars represent inter-replicate standard deviations. Log-rank-test.

To corroborate the critical relevance of the calcineurin/hsp90 axis for TSC-induced hype-rvirulence, we employed previously described *M. circinelloides* mutants harboring a loss-of-function of the two calcineurin catalytic A subunits (*cnaAΔ* and *cnaBΔ*) and regulatory B subunit (*cnbRΔ*) (33–34). Expectedly, an isogenic wild-type control (R7B) strain of *M. circinelloides* displayed significantly enhanced virulence after undergoing TSC, with 7-day survival rates of infected flies dropping from 65 % (static control) to 41 % (p = 0.004, Fig. 4E). In contrast, no significant difference in pathogenic capacity was seen between static and shear-challenged spores for all three *M. circinelloides* calcineurin loss-of-function mutants tested (Fig. 4E), thus providing further support for a model whereby TSC elicits transient hyper-virulence of Mucorales in a calcineurin-dependent way.

## Discussion

Clusters of NMM cases in patients suffering trauma in settings of extreme shear forces such as blast injuries or tornados gave rise to the hypothesis that extreme mechanical forces may impact the virulence of Mucorales. Mimicking TSC *in vitro* by high-speed cyclonical rotation on a magnetic stirrer, we found increased pathogenicity of Mucorales in our fruit fly infection model as documented by excess mortality of infected flies. Interestingly, other types of mechanical forces such as centrifugation or vortexing did not result in increased virulence, suggesting that the nature and intensity of physical forces determine differential effects on fungal biology. Accordingly, all NMM case patients after the deadly 2011 Joplim tornado had been located in the most severely damaged core of the tornado path (10). Previous work has established that the tangential flow and velocity fields in a magnetic stirrer are dynamically similar to that of big atmospheric vortices such as tornadoes (35). As the rotational speed and viscous effects in the rotating flow of a common laboratory stirrer are several magnitudes smaller than in a tornado (35), prolonged exposure time (≥ 30 minutes) was needed to induce Mucoralean hyper-virulence. Although most tornados last for only a few minutes, violent tornados such as the 2011 Joplin tornado can reach path lengths over 100 miles and travel for more than 30 minutes (36–37), underlining the physiological relevance of our experimental setting.

Unexpectedly, we found that Mucoralean hyper-virulence was not related to changes in fungal proliferation or capacity to cope with adverse conditions such as oxidative stress challenge, antifungal agents, or exposure to phagocytes. In line with these observations, RNA sequencing analysis of *R. arrhizus* revealed a very low number of significantly differentially expressed transcripts upon TSC. Specifically, no genes that are known or suspected to be immediately linked with Mucoralean virulence such as epithelial invasins, toxins, or proteins related to iron metabolism (38) were differentially expressed after TSC. While the identification of novel virulence factors and interpretation of the few differentially regulated transcripts is complicated by the sparse annotation of the *Rhizopus* genome and very limited experimental characterization of Mucoralean gene functions, the overall minimal transcriptional changes and rapid decay of hyper-virulence suggest a role of post-translational events that would have not been captured by our sequencing approach.

Our findings indicate that Mucoralean hyper-virulence after TSC is, to a large extent, driven by soluble metabolites that are released by shear-challenged spores and can subsequently increase the pathogenicity of static control spores. While struggling to solve the enigma of identifying the causative secreted factors, characterization of the overarching regulatory cascades driving shear stress-induced hyper-virulence could provide cues for potential therapeutic targets to improve the detrimental outcomes of NMM after high-energy trauma. Earlier studies of fungal homeostatic stress response following physical, oxidative, and antifungal challenge have revealed the evolutionary conserved calcineurin/hsp90 pathway as a fungal Achilles’ heel due to its role as a multifunctional regulator of cell wall integrity, adaption to adverse environments, and virulence (31–34, 39–40). Therefore, we hypothesized that this pathway may also be a gatekeeper of the Mucoralean response to TSC. Indeed, single and dual pharmacological inhibition of the calcineurin/hsp90 axis fully reverted TSC-induced hyper-virulence in *R. arrhizus* and the use of well-defined *M. circinelloides* calcineurin mutants (33–34) corroborated the pharmacological phenocopy. Calcineurin A has been shown to stress-dependently associate with endoplasmatic reticulum membranes in *Cryptococcus neoformans* and has been hypothesized to influence fungal membrane trafficking and protein folding (41). These observations constitute a potential role of calcineurin in the post-translational stage of protein biosynthesis that could possibly contribute to stress-induced alterations of the Mucoralean secretome, an area that would warrant in-depth exploration in future studies.

While calcineurin homologues are widely encountered throughout eukaryotic kingdoms including fungi (42), unusually high numbers of calcineurin pathway components were identified in Mucorales and have been implicated in Mucoralean virulence by governing dimorphic transitions and cell wall integrity (33–34, 39). Inhibition of the calcineurin pathway sensitized Mucorales to azole-induced apoptotic death and led to improved *in vitro* and *in vivo* efficacy of azole antifungals (43–45), highlighting its relevance to Mucoralean stress tolerance. Furthermore, recent reports have revealed non-canonical RNA interference pathways of *M. circinelloides* as post-transcriptional regulators of pathogenesis-related molecular processes including calcineurin pathway components in response to stressful stimuli (34, 46). This mechanism would deserve further characterization in the context of TSC-induced hyper-virulence. Importantly, the cited reports primarily described a role of the calcineurin/hsp90 pathway in the maintenance of cellular functionality and pathogenicity following exogenous stress events. In contrast, our observations in the TSC setting suggest that specific environmental stress stimuli can enhance Mucoralean virulence beyond the baseline level encountered under homeostatic conditions in a calcineurin/hsp90-dependent manner.

Our findings should encourage future studies of calcineurin inhibitor therapy in trauma-related NMM after high-energy events. Whereas systemically applied calcineurin inhibitors can potentiate immune paralysis, cause drug-drug interactions with many commonly applied antifungals (47), and exert adverse effects on wound healing (48), topical administration of calcineurin inhibitors to the wound environment could present an appealing alternative. Traditional antifungals perform poorly in NMM, as effective concentrations are difficult to achieve in inflamed or necrotic tissue (49) and the causative fungal agents are frequently multi-drug resistant (50). As calcineurin signaling is a cornerstone in the evolution of antifungal drug resistance, calcineurin inhibitors are less likely to induce resistance and could even mitigate resistance to co-administered topical or systemic antifungal therapy (40).

This study has some limitations. *Drosophila melanogaster* has been well-validated as a rapid, genetically amenable tool to study the virulence and immunopathogenesis of filamentous fungi (23–24) and there is robust evidence that Mucorales employ common virulence strategies to invade evolutionarily disparate organisms such as *Drosophila* and mammalians, highlighting the model’s suitability for primary comparative virulence screens (24). Nonetheless, confirmatory evidence in mammalian wound infection models (44, 51) would be warranted. While this was a pathogen-centered study, future investigations would also need to characterize the impact of TSC on fungal interactions with the diverse innate and adaptive immune cell repertoire of mammalian hosts, the local inflammatory environment in infected myocutaneous tissue, and biochemical parameters of wound healing. In addition, comparative studies in immunocompetent and immunosuppressed animals would be needed to account for trauma-induced immune paralysis (1, 52–53).

Furthermore, our study in a fungal mono-infection model cannot recapitulate the complexity of trauma-related soft tissue infections that are often polymicrobial in nature (1, 9–15). Cross-kingdom interactions of pathogens have been increasingly recognized as key virulence determinants shaping the outcomes of life-threatening infectious diseases (54–55). Direct physical interaction, inter-kingdom signaling, altered immunopathology, and competition for nutrients or trace elements are considered to play a driving role in the mutual modulation of bacterial and fungal virulence (54, 56). Interestingly, preliminary results of *R. arrhizus* and *Staphylococcus aureus* co-infection studies in our fruit fly model suggest that TSC-induced hyper-virulence rather increases in a mixed infection setting (Wurster et al., unpublished data), highlighting a need to obtain a more refined understanding of the influence of shear forces on the complex inter-dependencies in polymicrobial infections.

Despite these limitations, this study, for the first time, blends Mucoralean mechanobiology and pathogenicity and contributes to the understanding of NMM clusters after high-energy trauma events by revealing a hyper-virulent phenotype induced by tornadic shear stress. Furthermore, we identified an overarching pathway, whose pharmacological or genetic inhibition fully attenuated increased pathogenicity of shear-challenged Mucorales. This suggests new avenues of adjunct therapeutic interventions in order to improve the detrimental outcomes of NMM in trauma victims.

## Materials and Methods

### Fungal culture and shear stress exposure

The sources and culture conditions of fungal strains used in this study are summarized in **Table S1**. Spores were collected in saline by gently scraping the mycelium with a sterile glass rod. Fungal suspensions were washed twice with sterile saline and spore concentrations were determined using a hemocytometer. Fungal spores were diluted in 30 ml sterile PBS at a concentration of 1 □ 10^7^/ml. Depending on the experimental setup, 100 μg/ml cyclosporine A (CsA, Sigma-Aldrich), 50 μg/ml tanespimycin (17-N-allylamino-17-demethoxygeldanamycin, 17-AAG, Sigma-Aldrich), and/or 1 mM hydrogen-peroxide (H_2_O_2_, Fisher Scientific) were added. The highest non-inhibitory concentrations of 17- AAG and H_2_O_2_ were determined in preceding experiments using 2-fold serial dilutions. The spore suspension was stirred for at least 30 min in a 125 ml Erlenmeyer flask using a Corning PC-353 magnetic stirrer set to maximum speed (~ 1100 rpm). In pilot experiments to establish the optimal shear challenge procedure, spore suspensions were centrifuged at 6000 g for 30 min or shaken in a 50 ml tube taped to a vortex adaptor at maximum speed for 30 min.

### Drosophila melanogaster infection model

For Mucorales and *Fusarium* infections, female Oregon^R^ wild-type (WT) *Drosophila melanogaster* flies were used. As WT flies are resistant to *Aspergillus* infections (57), *A. fumigatus* was tested in female *Tl^r632^/Tl^I-RXA^ Drosophila* mutant flies, generated by crossing thermosensitive allele of *Toll (Tlr^632^)* flies with null allele of *Toll (Tl^I-RXA^)* flies. Standard procedures for the manipulation, housing, and feeding of flies were used as previously described (23–24). The dorsal side of the thorax of CO_2_-anesthetized flies (7-14 days old) was pricked with a size 000 insect pin (Austerlitz) dipped into the spore suspensions. Unless indicated otherwise in the figure legends, 1 □ 10^7^/ml spore suspensions were used and infections were performed within 10 min after termination of the stirring process. Flies were kept at 29 °C and transferred into fresh vials every other day. Survival was assessed daily until day 7 post-infection. At least three independent experiments with 20-26 flies per experiment and condition were performed on different days.

### Fungal RNA isolation

2 □ 10^7^ spores were mixed with 1 ml RNA*later* RNA stabilization reagent (Qiagen) and centrifuged at 17,000 g for 5 min. The supernatant was discarded and spores were resuspended in 500 μl buffer RLT (Qiagen) supplemented with 1 % β-mercaptoethanol. Bead-beating was performed for 2 □ 30 s using a Mini Beadbeater (Biospec Products) and UltraClean Microbial RNA Bead Tubes (Mo Bio), followed by a 3 min incubation step at 56 °C. Thereafter, RNA was isolated using the RNeasy Plant Mini Kit (Qiagen) according to the manufacturer’s instructions. RNA yield and purity were determined with a Nanodrop spectrophotometer (Thermo Scientific) and Agilent 2100 Bioanalyzer.

### RNA-sequencing and gene expression analysis

RNA-seq libraries (strand-specific, paired end) were generated from total fungal RNA using the TruSeq RNA sample prep kit (Illumina). Seventy-five nucleotides of the sequence were determined from both ends of each cDNA fragment using the HiSeq 4000 platform (Illumina). Sequencing reads were aligned to the reference *R. delemar* 99-880 genome using HISAT (58) and alignment files were used to generate read counts for each gene. Statistical analysis of differential gene expression was performed using the DEseq package from Bioconductor (59). A gene was considered differentially expressed if the FDR-value for differential expression was less than 0.05. The RNA-seq analysis was performed in biological triplicate.

### Statistics

Data analysis was performed using Microsoft Excel 2013 and GraphPad Prism 7.03. The log rank test (Mantel Cox test) was used to compare survival curves. For *in vitro* readouts, the two-sided t-test or one-way ANOVA with Tukey’s post-hoc test was used for significance testing depending on the data format. A p-value < 0.05 was considered significant.

## Author Contributions

DPK conceived and supervised the study. SW, AMT, VMB, and DPK designed experiments. SW, AMT, NDA, ACS, CMcC, KTG, and VMB performed experiments. SW, AMT, ACS, CMcC, VMB, and DPK analyzed data. ASI, JH, and SCL contributed fungal strains. ASI, JH, SCL, and AGM provided critical feedback. SW and DPK wrote the paper. All authors provided revisions and approved the final version of the manuscript prior to submission.

## Acknowledgments

DPK acknowledges the Texas 4000 Distinguished Professorship for Cancer Research and the NIH-NCI Cancer Center CORE Support grant no. 16672. ASI is supported by NIH/NIAID grant R01AI063503. JH is supported by NIH/NIAID MERIT Award R37-AI39115-21. He is a fellow and co-director of CIFAR program Fungal Kingdom: Threats & Opportunities. SCL is supported by the Max and Minnie Tomerlin Voelcker Fund. This study was also supported in part by NIH/NIAID grants U19AI110820 and R01AI141360 to VMB.

---

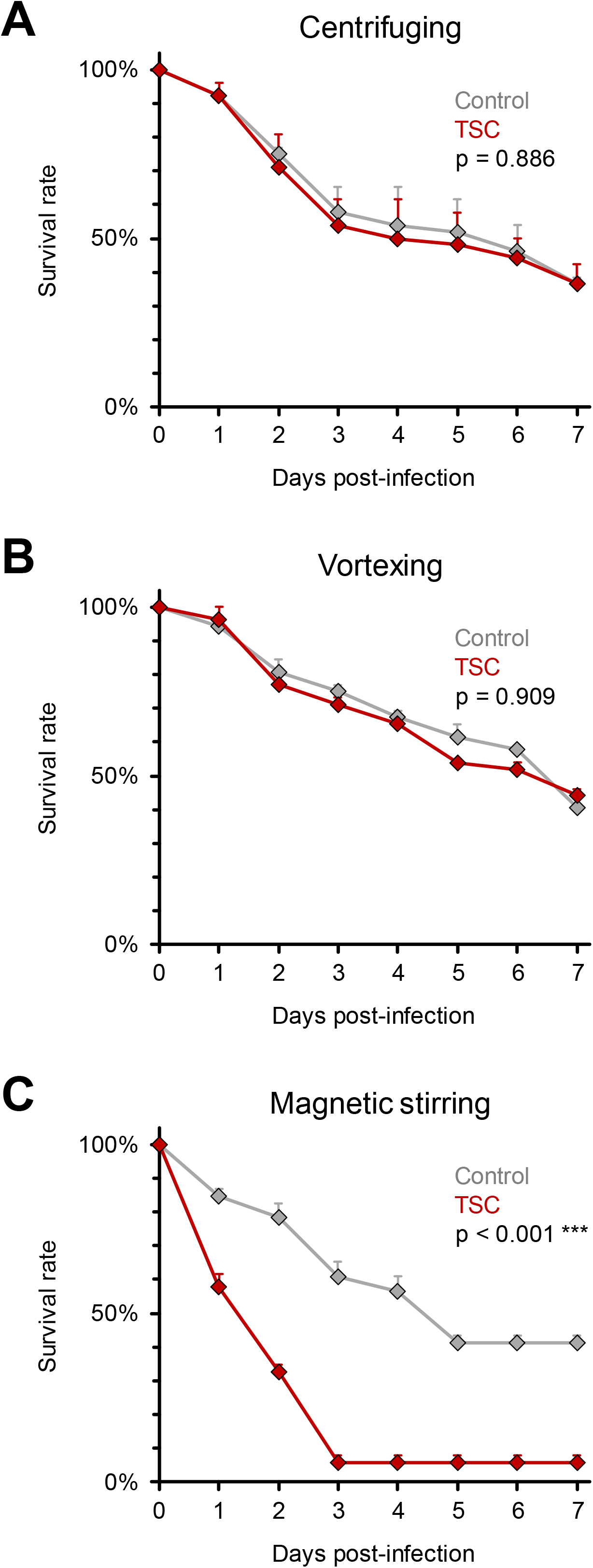

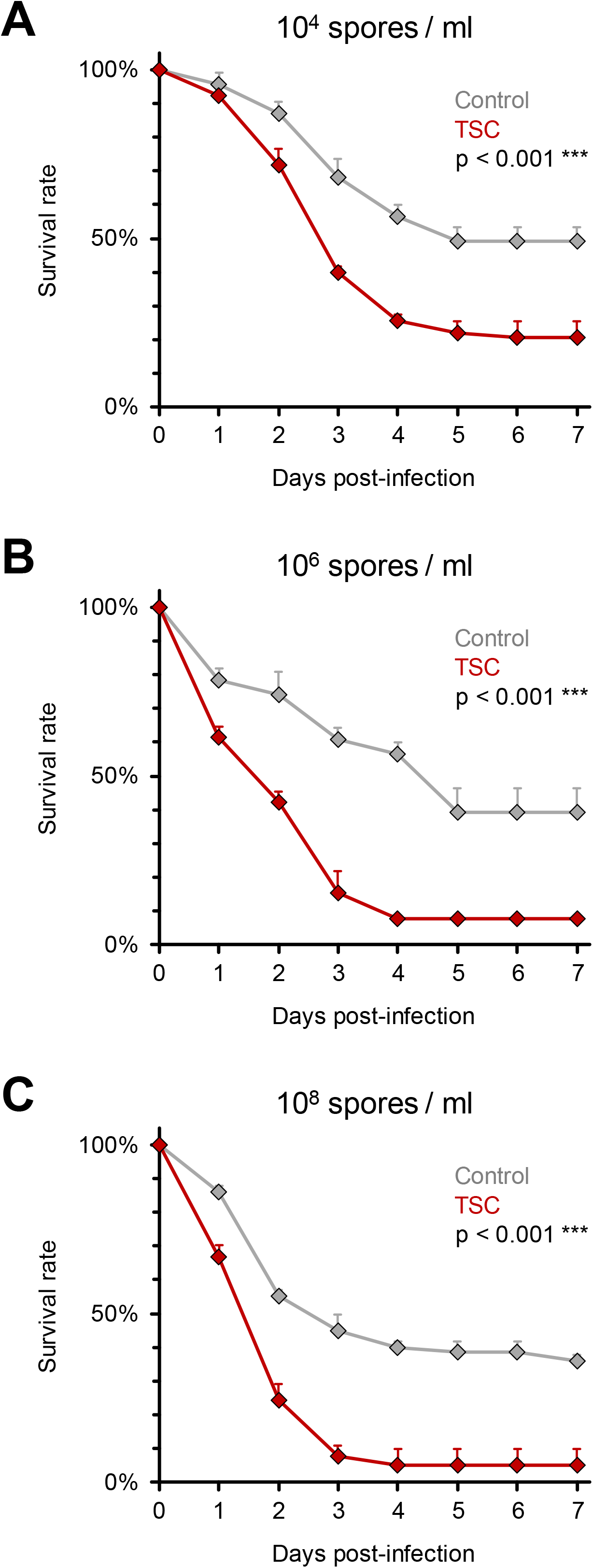

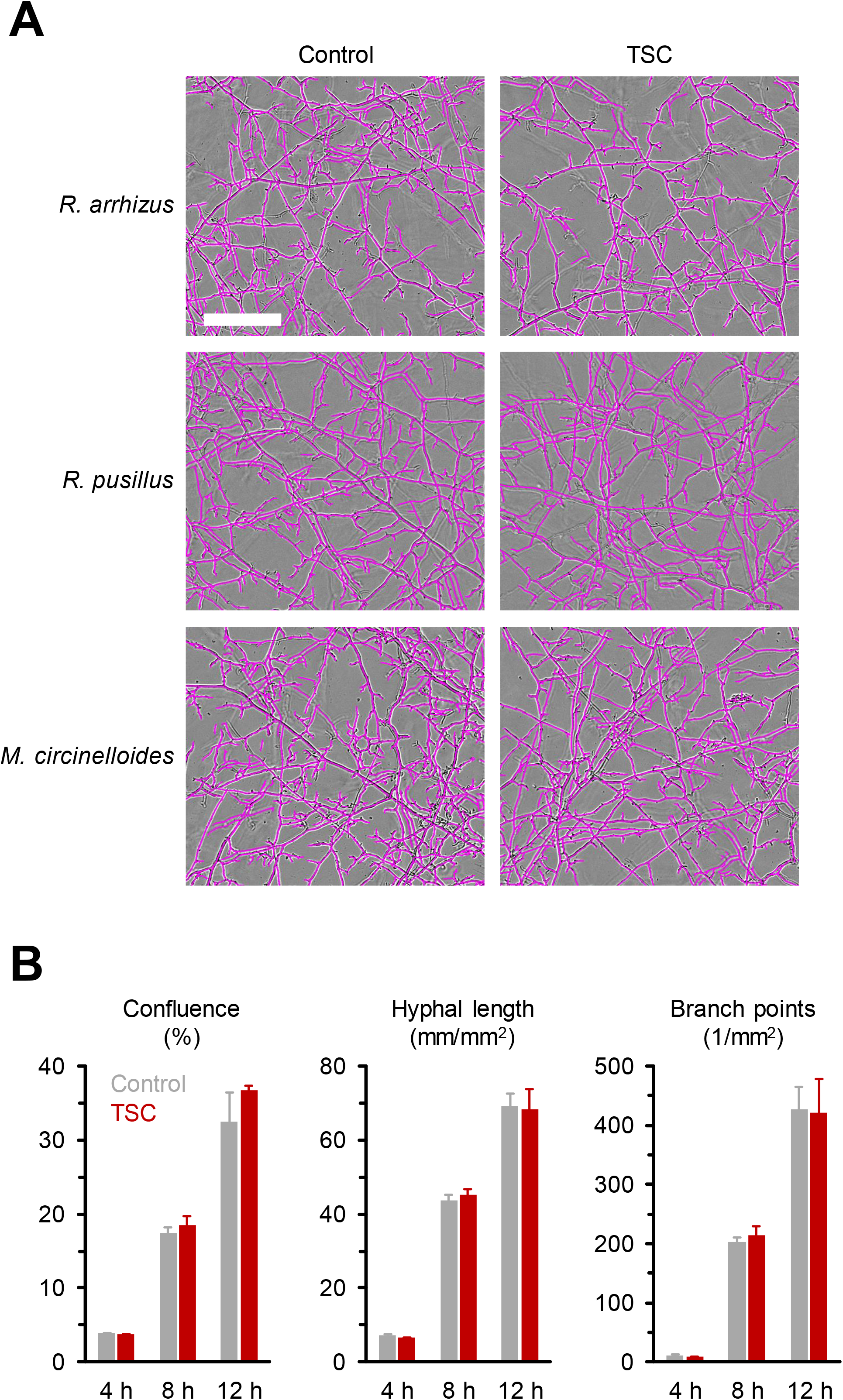

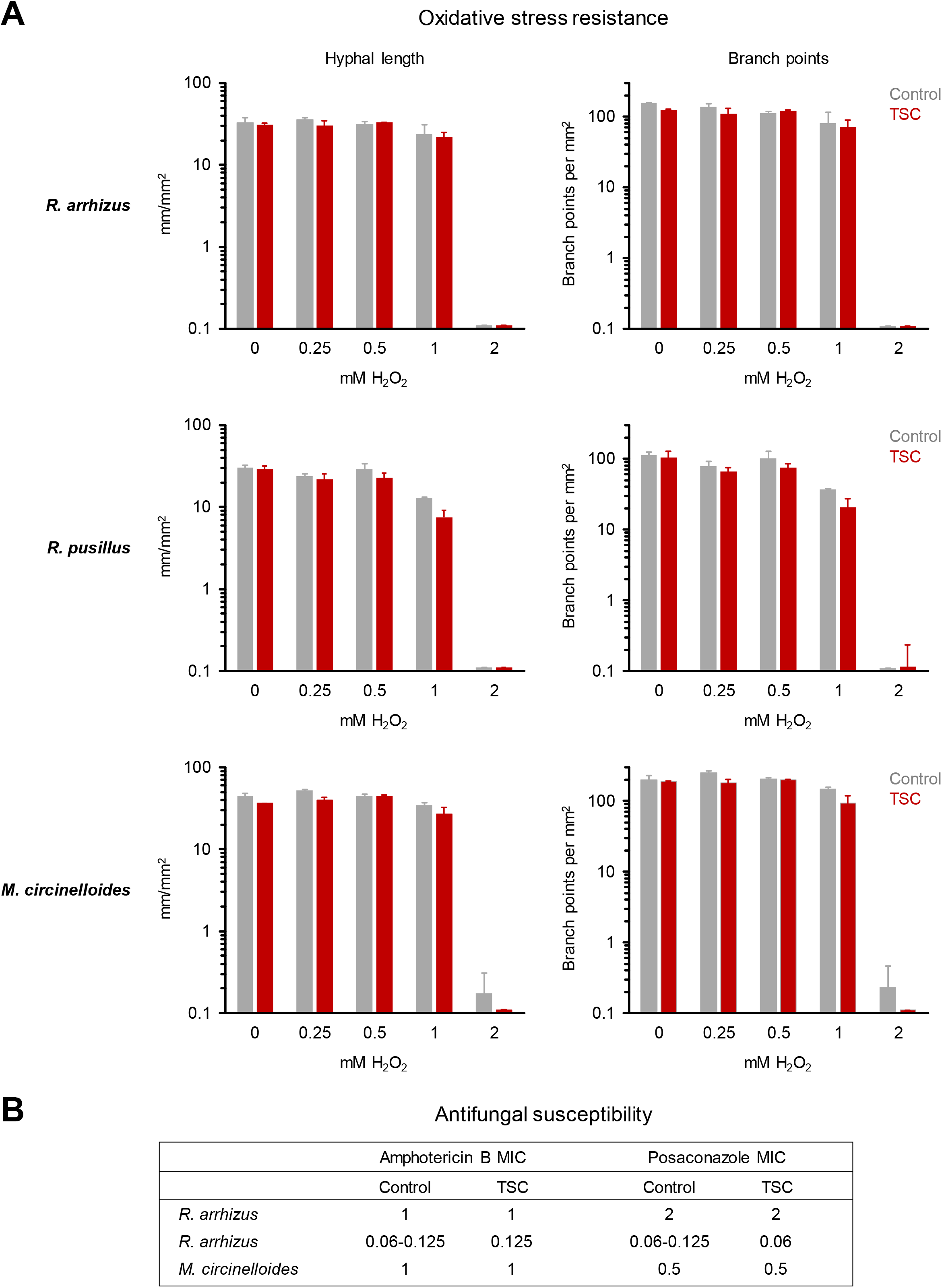

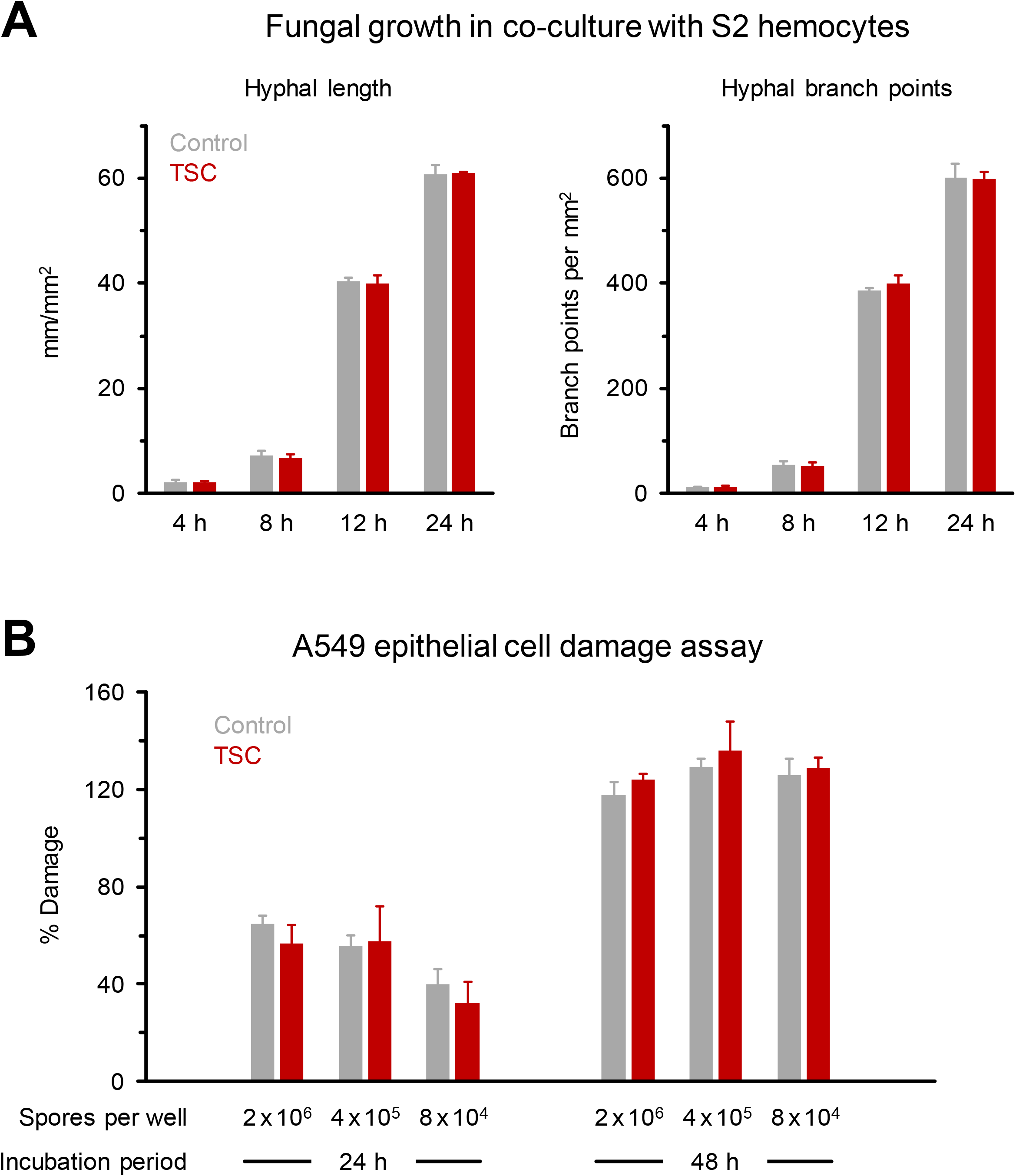

