## Supplementary Information for "Tornadic shear stress induces a transient, calcineurin-dependent hyper-virulent phenotype in Mucorales molds"

Supplementary Methods

*IncuCyte assay to monitor fungal morphology, mycelial expansion, and susceptibility to noxious*

*environments*

After termination of the magnetic stirring process to exert tornadic shear stress (TSC), spore suspensions were promptly diluted in RPMI + 2 % glucose at a concentration of 1 ᵡ 10^3^ spores per ml. 200 spores were seeded per well of a 96-well flat bottom plate. For selected experiments, serial dilutions of H_2_O_2_ (final concentration: 0.25 – 64 mM), amphotericin B (0.03-16 µg/ml), or posaconazole (0.03-16 µg/ml) were added. Phase images were obtained hourly for 24 h at 37 ˚C in the IncuCyte ZOOM HD/2CLR time lapse microscopy system (Sartorius) equipped with an IncuCyte ZOOM 10 ᵡ PLAN FLUOR objective (Sartorius). The IncuCyte image analysis software was used to quantify mycelial confluence, hyphal length, and branch point numbers as described before (1).

Hemocyte phagocytosis assay

*Drosophila* Schneider 2 (S2) cells (Gibco) were cultured in complete Schneider’s medium containing 10 % heat-inactivated fetal bovine (FBS, Sigma) serum, 0.1 % Pluronic F-68 (Gibco), and 50 IU/ml penicillin G + 50 μg/ml streptomycin sulfate (Gibco). Cells were kept in 75 cm^2^ culture flasks at 28 ˚C and passaged every 3-4 days as per the manufacturer’s recommendations. For co-culture experiments, S2 cells were centrifuged at 100 g for 5 min, quantified with a hemocytometer, and diluted in fresh complete Schneider’s medium to a concentration of 10^5^/ml. 100 µl aliquots (10^4^ cells) were combined with 10^3^ resting or shear-challenged *R. arrhizus* FTR1-GFP spores diluted in 100 µl complete Schneider’s medium in a 96-well flat bottom plate. The plate was imaged hourly (phase and green fluorescence, 400 ms acquisition time) in the IncuCyte ZOOM time-lapse microscopy system for 24 h at 28 ˚C. Hyphal length and branch point numbers per mm^2^ were quantified by NeuroTrack analysis as described before (1).

*Measurement of Rhizopus-induced host cell damage*

*Rhizopus*-induced A549 cell damage was quantified using the Pierce LDH Assay with slight modifications to the manufacturer’s protocol. Briefly, A549 cells were grown in 96-well tissue culture plates for 18 – 24 hours in F12k medium with L-glutamine plus 10 % FBS. Spores from *R. delemar* strain 99-880 (a clinical isolate obtained from a patient with rhinoorbital mucormycosis) were washed, resuspended in PBS, and then subjected to TSC by magnetic stirring or allowed to sit without stirring (control). Thereafter, spores were added to A549 cells at three different concentrations (2 ᵡ 10^6^, 4 ᵡ 10^5^, or 8 ᵡ 10^4^ spores per well). After 24 and 48 hours of incubation at 37 ˚C, 50 μl of the cell culture supernatant was collected from each well and transferred to a 96-well plate to assay for LDH activity. LDH is a cytosolic enzyme but will be released into the cell culture medium upon cell membrane damage. The amount of extracellular LDH is proportional to the amount of cell damage. Lysis buffer was added to all infected wells and incubated for 45 min at 37 ˚C. After lysis, 50 μl of cell culture supernatant was transferred to another 96-well plate and used for the LDH Assay Kit per protocol. LDH release was calculated as previously described (2).

Supplementary Figures

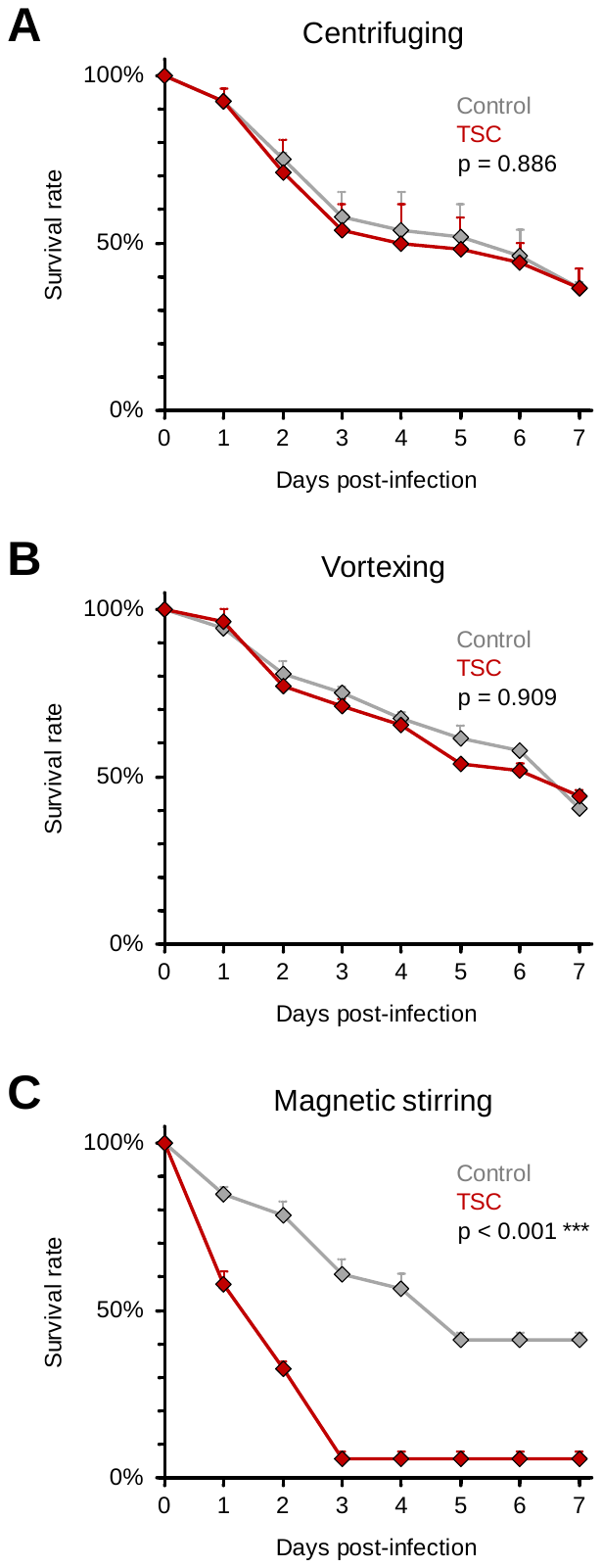

Fig. S1. Comparative evaluation of laboratory procedures for shear stress exposure of Mucorales spores.

*R. arrhizus* Ra-969 spore suspensions (10^7^/ml) were exposed to TSC by centrifugation at 6000 rpm (A), vortexing (B), or magnetic stirring at ~1100 rpm (C) for 30 min. Controls were kept in static culture for the same time. WT *D. melanogaster* flies were pricked with a needle dipped into the spore solutions. Two independent experiments were performed with a total of 46-52 flies per condition. Survival curves were compiled from aggregated results. Error bars represent inter-replicate standard deviations. Log-rank-test.

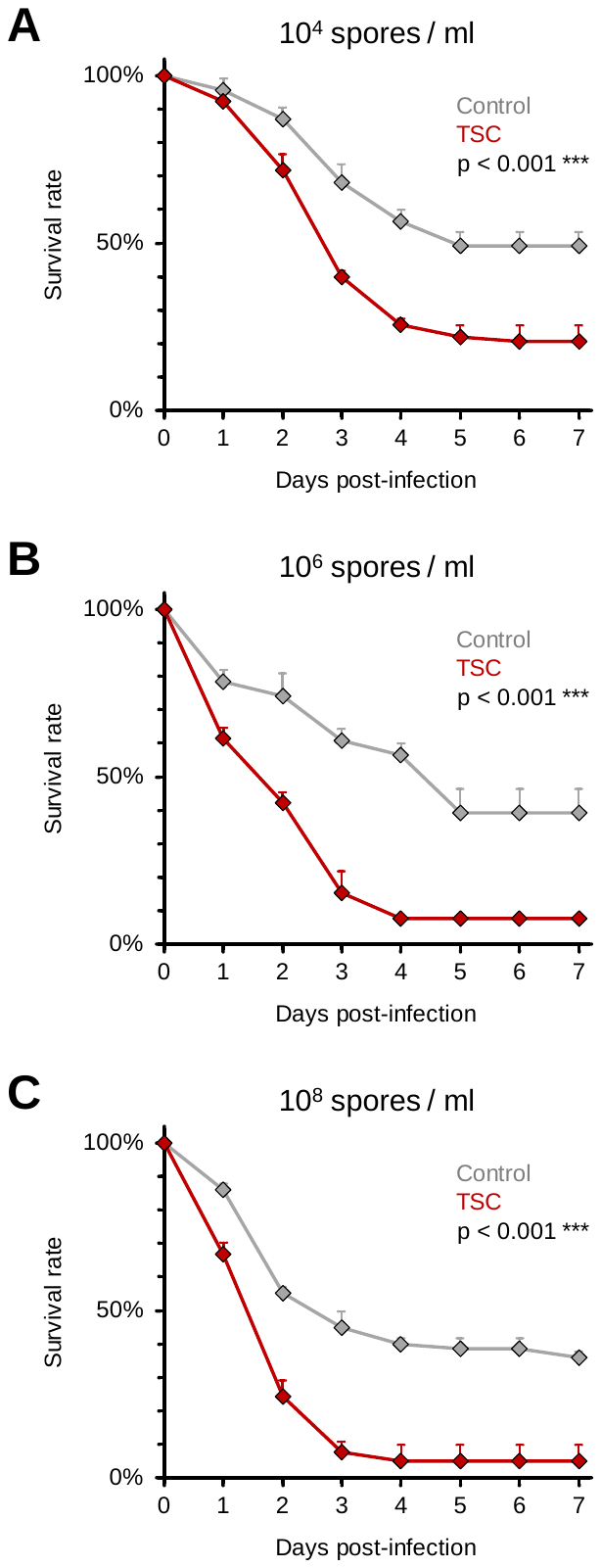

**Fig. S2.** **TSC-induced hyper-virulence is encountered across a wide range of spore concentrations**

*R. arrhizus* Ra-969 spore suspensions were prepared at concentrations of 10^4^ (A), 10^6^ (B), and 10^8^ (C) spores per ml. Spores were subjected to TSC by magnetic stirring or kept in static culture for the same time (Control). WT *D. melanogaster* flies were pricked with a needle dipped into the spore solutions. Three independent experiments were performed with a total of 69-78 flies per condition. Aggregated results were used for survival curves. Error bars represent inter-replicate standard deviations. Log-rank-test.

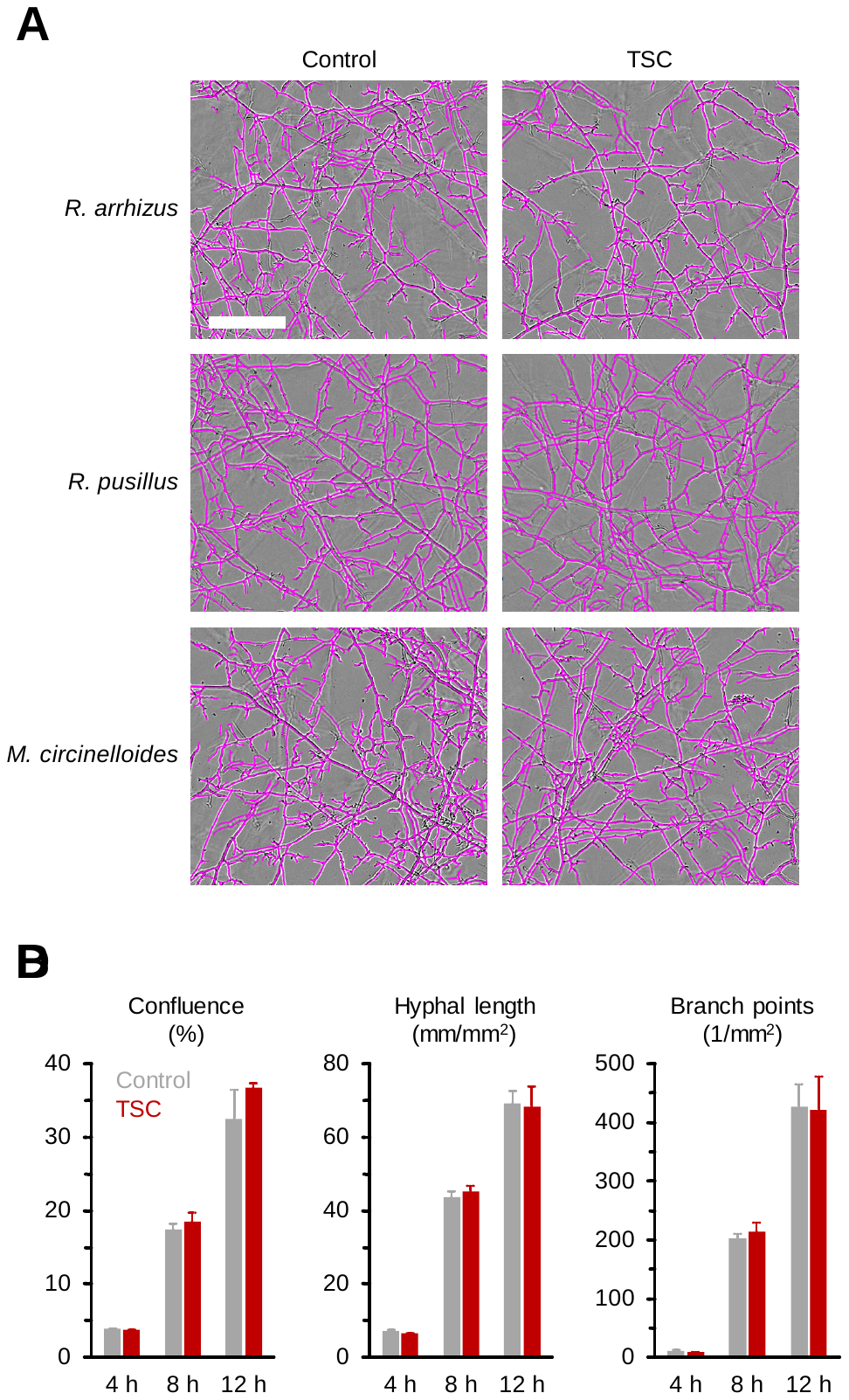

Fig. S3. TSC does not alter the morphology and mycelial expansion kinetics of Mucorales *in vitro*.

*Rhizopus arrhizus* Ra-749, *Rhizomucor pusillus* Rp-449, and *Mucor circinelloides* Mc-518 spore suspensions (10^7^/ml) were exposed to TSC by magnetic stirring for 30 min or kept in static culture (Control). Spore suspensions were subsequently diluted to 1 ᵡ 10^3^/ml in RPMI + 2 % glucose and 200 µl aliquots were added to 96-well flat bottom plates (200 spores per well). Plates were imaged hourly in the IncuCyte ZOOM time lapse microscopy system (37 ˚C). (A) Representative images after 12 h of culture are shown. Scale: 200 µm. (B) NeuroTrack (NT) and Basic Analyzer (BA) processing definitions were used to determine mycelial confluence (BA), total hyphal length (NT), and branch point numbers (NT) of Ra-749 after 4, 8, and 12 h of culture. Mean + SD (n = 3) are shown.

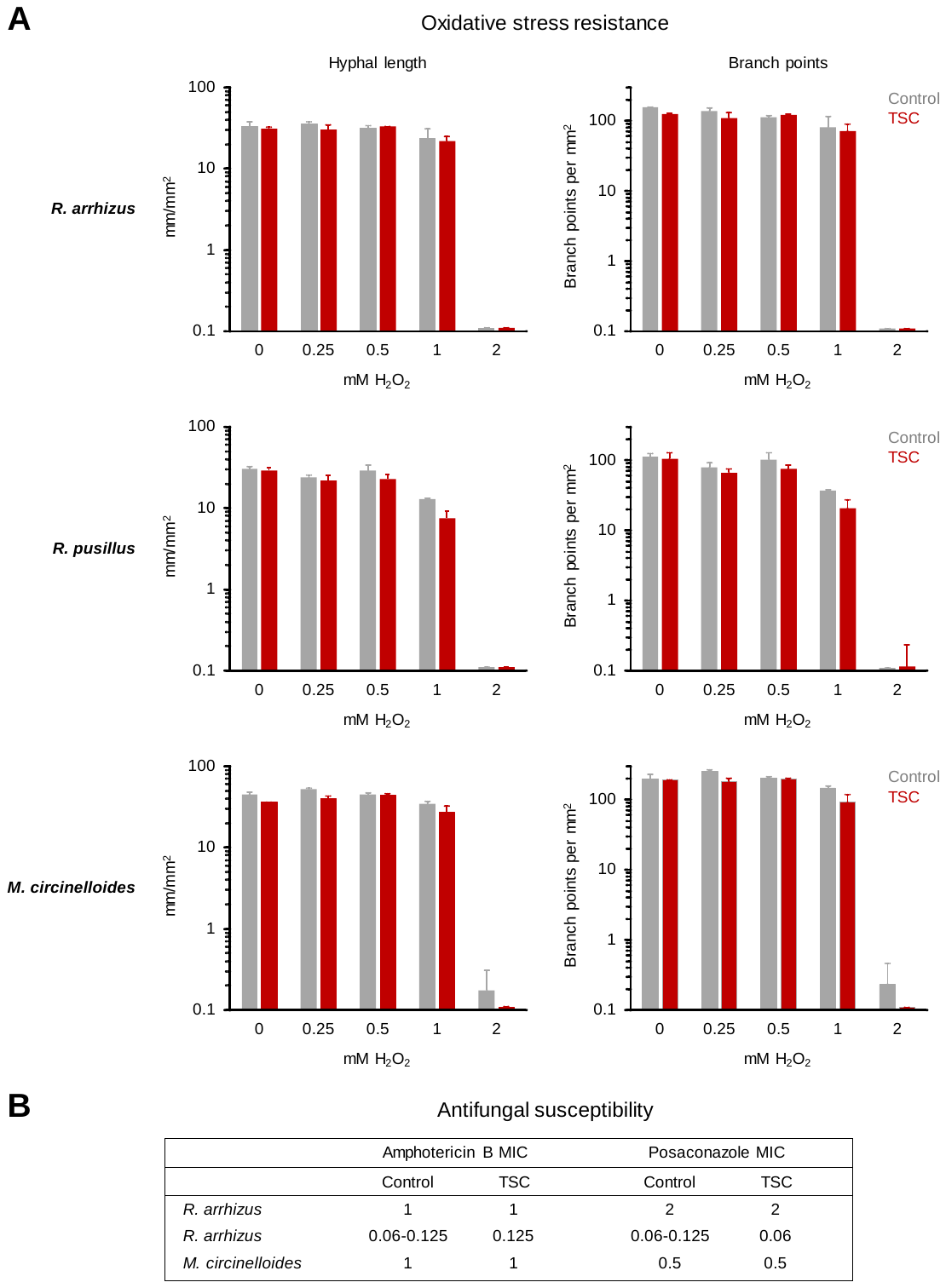

Fig. S4. The susceptibility of Mucorales to subsequent oxidative and antifungal stress remains unchanged after TSC

Legend on following page

Fig. S4. The susceptibility of Mucorales to subsequent oxidative and antifungal stress remains unchanged after TSC

*Rhizopus arrhizus* Ra-749, *Rhizomucor pusillus* Rp-449, and *Mucor circinelloides* Mc-518 spore suspensions (10^7^/ml) were exposed to TSC by magnetic stirring for 30 min or kept in static culture (Control). Spore suspensions were subsequently diluted to 2 ᵡ 10^3^/ml in RPMI + 2 % glucose. (A) 100 µl of the suspensions was added to 96-well flat bottom plates (200 spores per well) containing 100 µl RPMI with serial dilutions of H_2_O_2_ (final concentration, 0-32 mM). Phase images were obtained hourly for 24 hours in the IncuCyte ZOOM time lapse microscopy system. NeuroTrack processing definitions were used to determine hyphal length and branch point numbers depending on the H_2_O_2_ concentration and prior TSC exposure. Mean maximum hyphal length and branch point numbers (n = 4) observed during the 24 h observation period and standard deviations are shown. (B) 100 µl of the spore suspensions were mixed with serial dilutions of posaconazole (0.03-16 µg/ml) and amphotericin B (0.03-16 µg/ml) in 96-well round bottom plates and minimum inhibitory concentrations (MICs) were determined after 48 hours according to CLSI reference methods M38. Two independent replicates were performed.

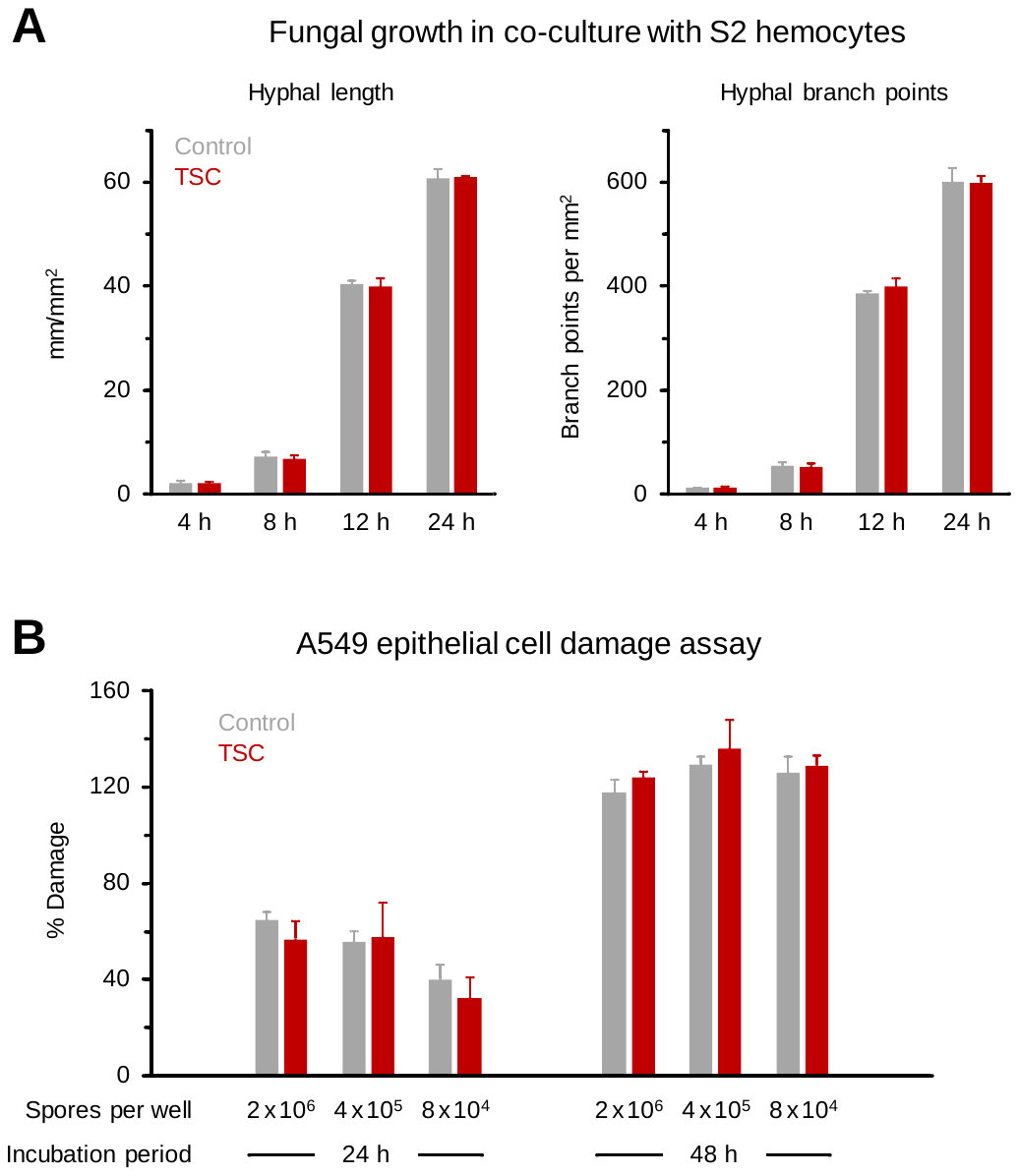

Fig. S5. Common virulence traits of Mucorales are unchanged after TSC

(A) Spore suspensions of GFP-expressing *R. arrhizus* (FTR1-GFP-*R. arrhizus,* 10^7^/ml) were either exposed to TSC for 30 min or kept under static conditions (Control), and diluted in complete Schneider’s medium at a concentration of 10^4^/ml. 100 µl aliquots of the suspensions (10^3^ spores) were combined with 10^4^ S2 hemocytes (effector-target ratio 10:1) diluted in 100 µl complete Schneider’s medium in a 96-well flat bottom plate. The plate was incubated in the IncuCyte ZOOM time-lapse microscopy system for 24 h at 28 ˚C. Phase and green fluorescence (400 ms acquisition time) images were obtained hourly. Mycelial length and branch points were quantified by NeuroTrack analysis. Mean + SD are shown (n = 3). (B) A549 alveolar epithelial cells were infected with *R. delemar* 99-880 spores that were subjected to TSC or kept under static conditions. A549 cell damage was quantified by LDH measurement as described above. Data are expressed as mean + SD based on three technical replicates.

Supplementary Table

Table S1. Fungal strains used in this study.

| **Species** | **Strain** | **Source/reference** | **Agar/medium** | **Temp.** | **Culture**  **duration** |
| --- | --- | --- | --- | --- | --- |
| *Rhizopus arrhizus* | Ra-749 | Clinical isolate ^a^ | Yeast extract agar | 37 ˚C | 2-3 days |
| *Rhizopus arrhizus* | Ra-969 | Clinical isolate ^a^ | Yeast extract agar | 37 ˚C | 2-3 days |
| *Rhizopus arrhizus* | FTR1-GFP-  *R. arrhizus* | (3) | Yeast nitrogen base agar + complete supplement mixture minus uracil | 37 ˚C | 3-5 days |
| *Rhizopus (arrhizus*  *var.) delemar* | RA 99-880 /  ATCC MYA-4621 | American Type  Culture Collection | Peptone dextrose agar | 37 ˚C | 3-5 days |
| *Rhizomucor pusillus* | Rp-449 | Clinical isolate ^a^ | Yeast extract agar | 37 ˚C | 2-3 days |
| *Mucor circinelloides* | Mc-518 | Clinical isolate ^a^ | Yeast extract agar | 30 ˚C | 2-3 days |
| *Mucor circinelloides* | R7B | (4-5) | Yeast peptone glucose agar | 30 ˚C | 5 days |
| *Mucor circinelloides* | *cnaAΔ* (MSL9) | (4-5) | Yeast peptone glucose agar | 30 ˚C | 5 days |
| *Mucor circinelloides* | *cnaBΔ* (MSL22) | (5) | Yeast peptone glucose agar | 30 ˚C | 5 days |
| *Mucor circinelloides* | *cnbRΔ* (MSL8) | (4-5) | Yeast dextrose agar | 30 ˚C | 2 days |
| *Apophysomyces*  *trapeziformis* | CBS 125534 | CBS-KNAW Fungal Biodiversity Centre | Minimal medium | 37 ˚C | 5 days |
| *Aspergillus fumigatus* | ATCC MYA-4609  (Af-293) | American Type  Culture Collection | Yeast extract agar | 37 ˚C | 2-3 days |
| *Fusarium solani* | Fs-001 | Clinical isolate ^a^ | Yeast peptone glucose liquid medium | 30 ˚C | 3-5 days |

^a^ Isolates were obtained from cancer patients at the University of Texas M.D. Anderson Cancer Center, Houston, Texas, USA.

**Supplementary Information References**
